# Volitional hand activation intensifies cortical proprioceptive processing in the primary sensorimotor cortex

**DOI:** 10.64898/2026.06.08.730851

**Authors:** Maija Siltala, Timo Nurmi, Maria Hakonen, Harri Piitulainen

## Abstract

Proprioception refers to the sense of position and movement of the body and is essential for the planning and execution of movements, especially skilled hand movements. The somatotopic organization of the fingers in the human sensorimotor (SM1) cortex has previously been studied using passive tactile stimulation and active movements. However, the finger representations in response to proprioceptive stimulation in the SM1 cortex are incompletely understood. To address this, we investigated cortical representations of the index and ring fingers during kinematically matched passive and active finger movements using 3T fMRI and an MRI-compatible proprioceptive stimulator. Both active and passive conditions elicited significant activations in the SM1 cortex, with stronger and more extensive responses observed during the active condition. The results indicated that volitional activation of the fingers intensifies cortical proprioceptive processing in the SM1 cortex. This observation could be explained by the active state of the SM1 cortex and the increased sensitivity of peripheral proprioceptors during the active condition.

**Key points:** Volitional activation of the fingers intensifies cortical processing of proprioceptive afference and modulates the organization of finger representations in the SM1 cortex.

This could be explained by the active state of the SM1 cortex and the increased sensitivity of peripheral proprioceptors during the active condition.

Proprioceptive finger representations followed the classical somatotopic organization at the group level, although this organization was not consistently observed across all subjects.

## 1 Introduction

The human primary sensorimotor (SM1) cortex exhibits a classical somatotopic organization, in which individual fingers are represented in distinct cortical regions (Penfield & Boldrey, 1937). This organization has also been studied using functional magnetic resonance imaging (fMRI). In the primary somatosensory (SI) cortex, finger representations in response to passive cutaneous tactile/vibrotactile stimulation and active movements have been mapped in multiple studies, consistently showing a lateral-to-medial and inferior-to-posterior organization from digit 1 (thumb) to digit 5 (little finger), located primarily in Brodmann areas (BAs) 3b, 1, and 2 (Janko et al., 2022). Similarly, in the primary motor (M1) cortex, finger representations in response to active finger movements have been shown to follow a somatotopic organization in which digits are arranged lateral-to-medial from digit 1 to digit 5 (Schellekens et al., 2018; Siero et al., 2014). However, the organization within the M1 cortex has been shown to be more complex than the classical somatotopic organization at a finer spatial scale. Individual digits can appear in multiple, mirrored representations that are differentially engaged depending on the specific motor action (grasping versus retraction) (Huber et al., 2020). Previous research has also shown that tactile stimulation and voluntary finger movements follow a somatotopic organization with a similar spatial pattern in the SI cortex (Sanders et al., 2023). However, voluntary movement has been shown to produce greater overall BOLD activity compared to tactile stimulation (Berlot et al., 2019; Sanders et al., 2023).

Proprioception, the sense of position and movement of the body, is critical for hand and finger movements (Long et al., 2022; Lutz & Bensmaia, 2021). In contrast to tactile and motor finger representations, cortical representations of proprioceptive afference to the human SM1 cortex are incompletely understood. Previous fMRI studies using proprioceptive stimulation have shown that passive movement elicits significant activation in the SM1 cortex, but these studies have focused on either single-finger movements or the simultaneous movement of multiple fingers (Blatow et al., 2011; Boscolo Galazzo et al., 2014; Lolli et al., 2019; Nasrallah et al., 2019; Reddy et al., 2001; Thickbroom et al., 2003). In addition, a recent 7T fMRI study investigated the laminar response of the human M1 cortex to proprioceptive stimulation of a single finger and found that passive movement activates both superficial and deep layers of the M1 cortex (Knudsen et al., 2025). However, it remains unknown how proprioceptive representations of multiple fingers are organized in both the M1 and SI cortices, and whether volitional muscle activation during proprioceptive stimulation modulates the finger representations.

The aim of this study was to investigate cortical representations of the index and ring fingers in the SM1 cortex during passive and resisted (active) finger movements using 3T fMRI and an MRI-compatible proprioceptive stimulator. Specifically, we clarified whether the index and ring fingers of the same hand show spatially distinct representations to proprioceptive stimulation in the SM1 cortex, and whether SM1 cortex activation differs in magnitude and spatial features between the passive and active conditions. Passive finger movements provide a natural way to stimulate peripheral proprioceptors, while the use of proprioceptive stimulator ensures consistent stimulation frequency and amplitude across participants.

## 2 Materials and methods

### 2.1 Participants

Twenty-four healthy participants (mean age ± standard deviation = 33.4 ± 7.3 years) with no neuropsychiatric diseases and no contraindications for MRI were recruited for the study. According to the Edinburgh Handedness Inventory (Oldfield, 1971), all participants except one were right-handed; one participant was ambidextrous (mean handedness score ± standard deviation = 89.4 ± 16.0). Six participants were excluded due to excessive head motion (see 2.4 Data preprocessing), leaving 18 participants in the final analysis. The study was approved by the ethics committee of the University of Jyväskylä and written consent was obtained from all participants prior to the study. The study was performed in accordance with the principles of the Declaration of Helsinki.

### 2.2 Experimental design

During MRI, participants lay in a supine position in the scanner, with their right fingers attached to a custom-made, MRI-compatible four-finger proprioceptive stimulator (University of Jyväskylä, Finland) fixed to the scanner table (Fig. 1a). Each ring-shaped finger holder was attached to a separate 20-cm long pneumatic muscle (DMSP-10-200 AM-CM, diameter 10 mm, Festo AG & Co., Esslingen, Germany) able to evoke ∼2 cm flexion-extension movement of the proximal interphalangeal joint of the fingers. The size of the finger holder was individually selected to prevent the finger from slipping out during movement. In addition, participants wore a soft glove to reduce tactile stimulation caused by contact between the fingers and the stimulator. An elastic band was wrapped around the participant’s right thumb to prevent it from touching the index finger. The index and ring finger movements were controlled using the Presentation software (Version 24.0, Neurobehavioral Systems, Inc., Berkeley, CA). Participants were instructed to remain still and avoid head movements during scanning.

**Figure 1.**
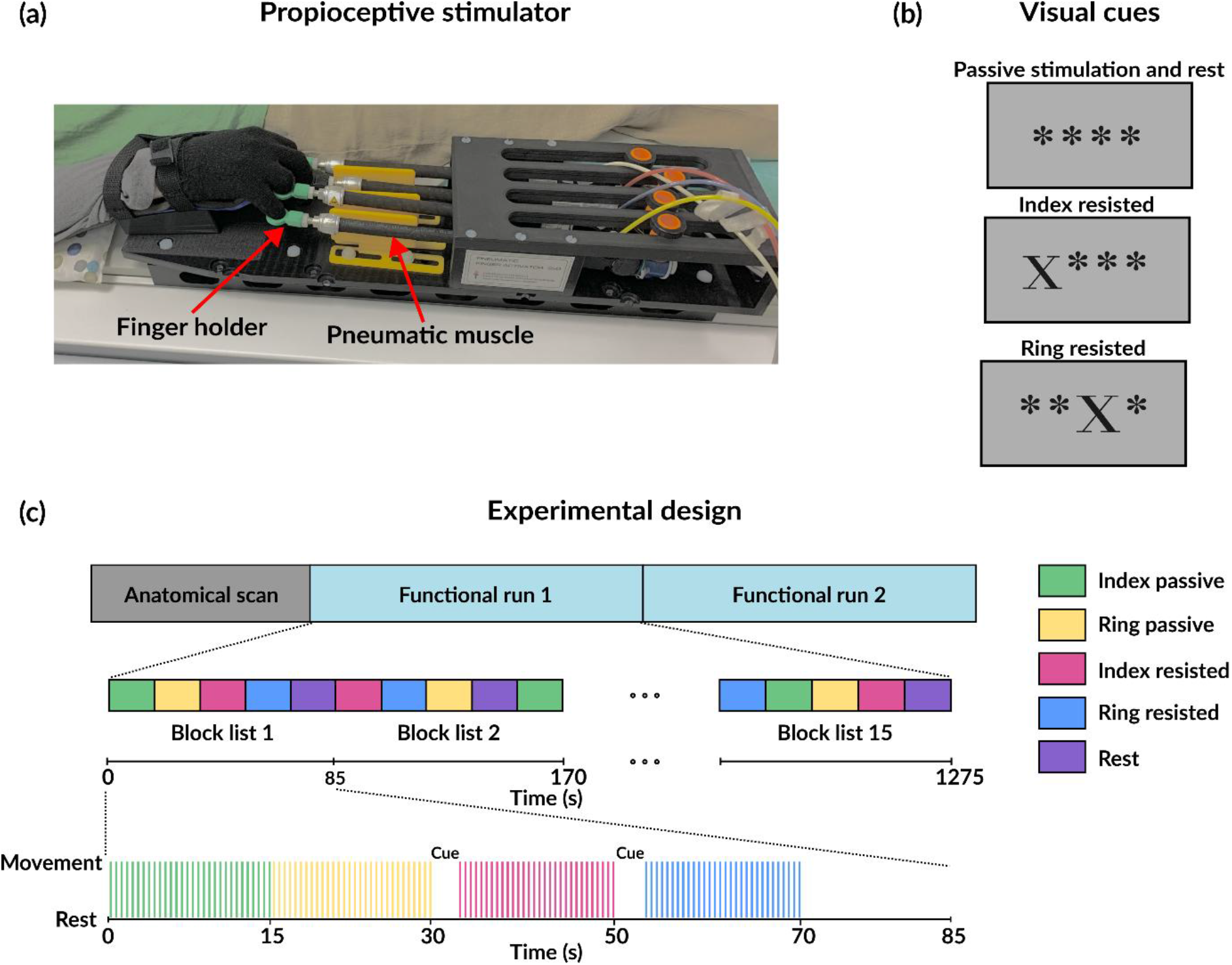
Proprioceptive stimulator and experimental design. (a) Participant lying on the scanner table with the fingers of her right hand attached to the proprioceptive stimulator. (b) Visual cues displayed during different conditions in the functional runs. (c) The imaging session included an anatomical scan and two functional runs. Each functional run consisted of a pseudorandomized block design with 75 condition blocks arranged into 15 block lists. Each condition appeared once per block list, and finger movement conditions included 30 individual movement cycles at 2 Hz. In active conditions, a 5-s cue was presented before the movement stimulation started.

Participants attended a single imaging session, which included an anatomical scan and two functional runs, each lasting 21.8 minutes. Each functional run consisted of a pseudo-randomized block deign with 15 blocks per condition. The five conditions were: 1) passive stimulation of the index finger, 2) passive stimulation of the ring finger, 3) resisted (active) stimulation of the index finger, 4) resisted (active) stimulation of the ring finger, and 5) rest. During passive stimulation, the stimulator moved the relaxed index or ring finger at 2 Hz which lies in the frequency range of normal hand movements, allowing smooth movement. In the resisted stimulation, the movement was the same, but the participant resisted the movement by applying a slight finger flexion. In the rest condition, the participant remained at rest, and the fingers were not moved. The blocks occurred in block lists of five, such that each condition was presented once per block list (Fig. 1c). During passive stimulation and rest, the participant was shown a screen with four stars, each representing one finger in the stimulator (Fig. 1b). During resisted stimulation, the star corresponding to the moved finger changed to an ‘X’ (Fig. 1b). The passive stimulation and rest blocks each lasted 15 s. At the beginning of each resisted stimulation block, a 5-s cue was first presented, and the participants started the active finger flexion at the onset of the 15-s movement stimulation, resulting in a total block duration of 20 s. The cue image was identical to the one used during the movement phase. In each block that involved finger movement, 30 trials were presented, with each trial corresponding to a single finger movement at 2 Hz.

### 2.3 Data acquisition

Imaging was performed using a Siemens 3T MAGNETOM Vida MRI scanner with a 64-channel receiving head coil. To minimize participant head movement during scanning, padding was placed around the head. Anatomical T_1_-weighted images were acquired with a magnetization-prepared rapid gradient echo (MPRAGE) sequence and the following parameters: TR = 2000 ms, TE = 2.41 ms, flip angle = 9°, resolution = 0.7×0.7×0.9 mm^3^, FOV = 216×230×187.2 mm^3^, orientation = axial oblique. The anatomical scan was followed by two functional runs, each comprising 1300 whole brain volumes. fMRI data were acquired with a simultaneous multi-slice (SMS) echo planar imaging (EPI) sequence and the following parameters: TR = 999 ms, TE = 31 ms, flip angle = 52°, resolution = 2.4×2.4×2.4 mm^3^, FOV = 216×216×144 mm^3^, orientation = axial oblique.

### 2.4 Data preprocessing

Anatomical and functional data were preprocessed using FreeSurfer 7.4.1 (Fischl, 2012). First, anatomical T_1_-weighted images were used to generate cortical reconstructions for each participant using the recon-all function. The reconstructed surface was evaluated for quality assurance, and segmentation errors in the pial and white matter surfaces were manually corrected with Freeview’s Recon Edit tool. The functional data were preprocessed using the FS-FAST pipeline within FreeSurfer. Volumes acquired after the functional task had ended were removed from each functional run. Preprocessing included motion correction, co-registration of the functional data to each subject’s anatomical image, intensity normalization, and spatial smoothing on the cortical surface using a 5-mm full-width at half-maximum (FWHM) Gaussian kernel. All individual-level analyses were performed in each subject’s native surface space, except for the calculation of geodesic distances between peak finger activation values and the assessment of whether activation patterns followed a somatotopic organization. For these exceptions, as well as for group-level analyses, the data were resampled to the fsaverage template (163842 vertices per hemisphere) (Fischl et al., 1999). The motion parameters estimated during the motion correction were used to assess the head movement. Data from six participants were excluded from further analysis due to excessive head motion. The exclusion criterion was defined as a vector translation exceeding 1.5 mm relative to the middle volume of the functional run at any time point and the presence of sudden peaks in head motion. Thus, the total number of participants included in the analysis was 18.

### 2.5 Data analysis

Figure 2 shows regions of interest (ROIs) that were anatomically defined for each subject in their native surface space and on the fsaverage template using FreeSurfer’s probabilistic Brodmann area parcellation. The SI cortex ROI was defined by combining thresholded BAs 1, 2, 3a, and 3b while the M1 cortex ROI was defined by combining thresholded BAs 4a and 4p. To focus the analysis on the hand region, both the SI and M1 cortex ROIs were subdivided into seven components using principal component analysis (PCA), with the subdivisions oriented orthogonally to the first principal component (Cammoun et al., 2012; Gramfort et al., 2013, 2014). From these components, subject-specific ROIs representing the SI and M1 cortex hand areas were constructed.

**Figure 2.**
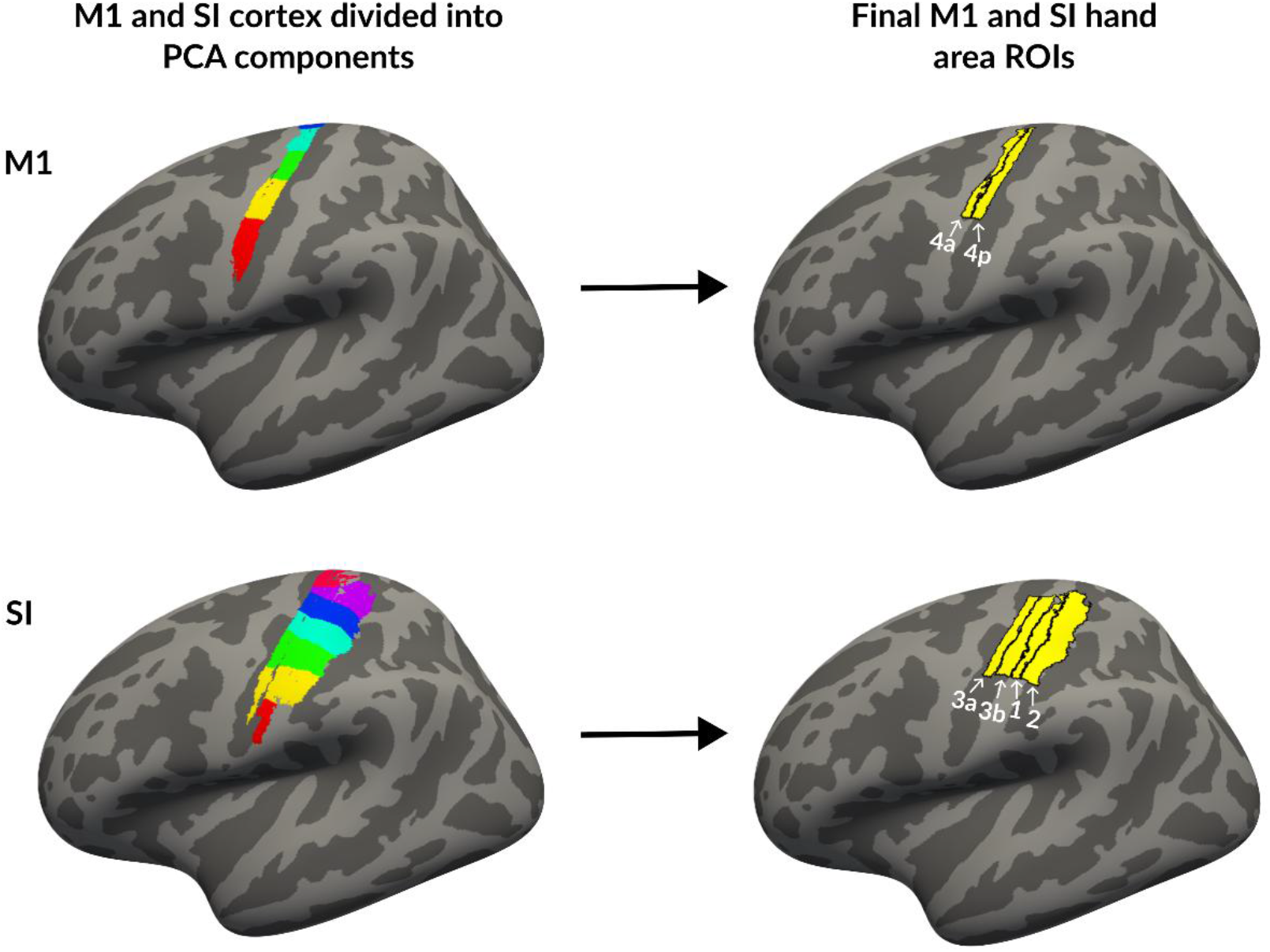
Determination of ROIs. On the left, M1 and SI ROIs are shown as defined by combining thresholded BAs 4a and 4p for the M1 cortex, and BAs 1, 2, 3a, and 3b for the SI cortex. These ROIs were further subdivided into seven components using PCA, with each component displayed in a different color. On the right, the final hand area ROIs are shown, constructed by selecting the same PCA components for each participant.

The first-level analysis was conducted using standard functions from FreeSurfer’s FS-FAST pipeline to fit a general linear model (GLM) to each cortical vertex. Analyses were performed in both the native surface space and the fsaverage template space. The hemodynamic response function (HRF) was modeled using the canonical SPM HRF function and its temporal derivative, and both were convolved with the task regressors prior to model fitting. Motion-related nuisance regressors derived from motion correction parameters (three translations and three rotations) were included in the GLM. For analyses conducted in native surface space, five contrasts were computed: each finger (active and passive) versus rest, and all conditions versus rest. For analyses conducted in fsaverage space, eight contrasts were computed: each finger (active and passive) versus rest, as well as active versus passive and passive versus active contrasts for both fingers. For individual-level analyses, the resulting activation maps were corrected for multiple comparisons using false discovery rate (FDR) (Genovese et al., 2002) at p < 0.05, applied within the subject-specific ROIs corresponding to the SI and M1 cortex hand areas. Group-level analysis was conducted using the FS-FAST pipeline to identify vertices for which contrast effects were consistent across participants. The resulting activation maps were corrected for multiple comparisons using cached Monte Carlo simulations with a voxel-wise threshold of p < 0.001 for positive contrast values and cluster-wise threshold of p < 0.05.

At the individual level, we compared the extent and strength of the activations between active and passive conditions for each finger in the M1 and SI cortices. To further characterize activation strength, we computed a mean percent signal change profile over the M1 and SI cortex Brodmann areas. Furthermore, the overlap and geodesic distance were compared between fingers within the passive and active conditions. Finally, it was examined whether the finger representations followed a somatotopic organization. The group-level activation maps were calculated only for visualization purposes. In addition to FreeSurfer, custom MATLAB (R2024b, Mathworks, Natick, MA, United States) and Python scripts were used to perform data analysis.

The extent of activation was calculated from finger versus rest significance maps, masked to the SI and M1 cortex hand areas. Extent was defined as the number of statistically significant vertices and was compared between active and passive conditions for each finger and ROI. For the strength of activation, a mask was created by contrasting all movement conditions against the rest. This mask was applied to the percent signal change maps, and the average percent signal change was computed. Strength values were then compared between active and passive conditions for each finger and ROI. The activation strength profile across the Brodmann areas of the M1 and SI cortices was obtained by dividing M1 ROI into 8 and SI ROI into 16 smaller bins (Supplementary figure 3), computing the mean signal change within each bin, and averaging these profiles across participants.

To assess overlap between finger activations, a modified Dice coefficient between fingers was calculated for passive and active conditions across both ROIs

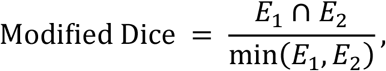

where *E*_1_ > 0 is the extent of index finger activation and *E*_2_ > 0 is the extent of ring finger activation as defined before. The modified Dice coefficient accounts for cases where one representation is fully contained within another but differs greatly in size (Sanders et al., 2023). The coefficient ranges from 0 to 1, with 0 indicating no overlap and 1 indicating complete overlap. Overlap was compared between active and passive conditions for both ROIs. The finger separation was quantified by computing the geodesic distance between the vertices showing maximum percent signal change for each finger. The geodesic distance was calculated using the Python package gdist (https://github.com/the-virtual-brain/tvb-gdist). These maximum values were obtained from the percent signal change maps masked to the M1 and SI cortex hand areas. Distances were compared between active and passive conditions for both ROIs.

Finally, we tested whether the finger representations followed somatotopy by calculating the average MNI x-coordinate from the finger-versus-rest maps. If the representations followed the somatotopic organization, the index finger would be located more laterally, corresponding to a more negative MNI x-coordinate. Statistical analyses were performed in MATLAB using the Wilcoxon signed-rank test, due to the small sample size and unequal number of participants across analyses. The resulting p-values were corrected for multiple comparisons using the Benjamini-Hochberg procedure to control the false discovery rate (Groppe, 2025).

## 3 Results

### 3.1 Activation extent and strength

Figures 3a and 3b show the comparison of activation extent and strength between the active and passive conditions for each finger in the M1 and SI cortices. Participants with no significant vertices in either the active or passive condition were excluded from the analysis (N = 12–18 depending on the variable analyzed). The extent of activation, defined as the number of statistically significant vertices, was larger in the active condition (resisted stimulation) compared to the passive condition for both fingers in the M1 and SI cortices (Supplementary table 1). To assess differences in the activation strength, we compared the average percent signal change between the active and passive conditions within the mask defined by all movement conditions versus rest. The strength of the activation was higher in the active condition compared to the passive condition for both fingers in the M1 and SI cortices (Supplementary table 1).

**Figure 3.**
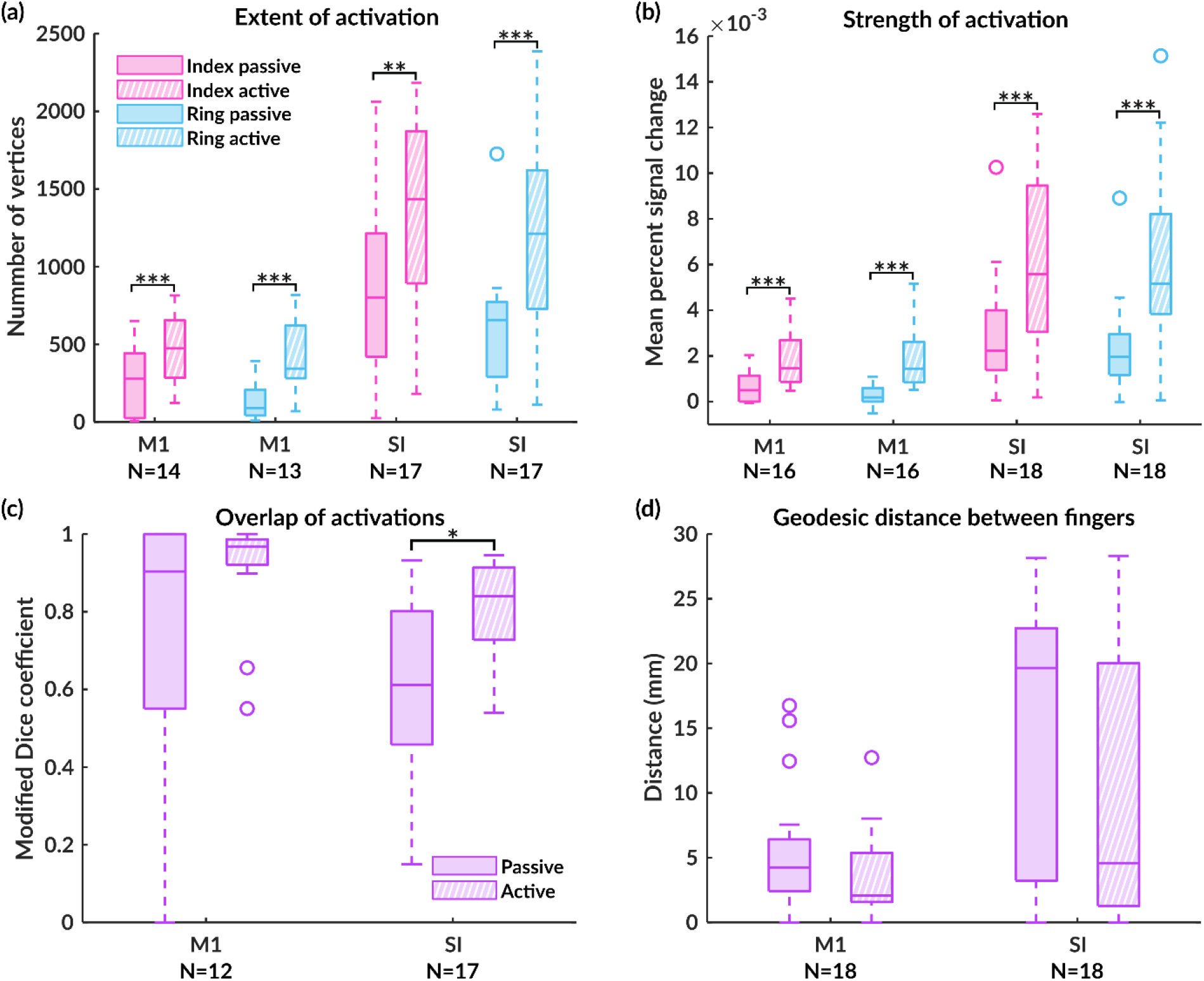
Metrics of cortical finger representations in the M1 and SI cortices during active and passive conditions. (a) Extent of activation, (b) Strength of activation, (c) Overlap between the fingers, and (d) Geodesic distance between the fingers. Significance is indicated by: *** = p < 0.001, ** = p < 0.01, and * = p < 0.05.

Figure 4a shows the group-average percentual signal change profiles across Brodmann areas in the M1 and SI cortices. Based on visual inspection, the overall shapes of the profiles were similar for both fingers in both active and passive conditions, although the passive condition consistently showed lower signal changes. In the M1 cortex, the profile peaked in BA 4p, near the approximate boundary between BA 4p and BA 3a. In SI, a higher peak was observed in BA 3a, near the approximate boundary between BA 3a and BA 3b, with an additional smaller peak in BA 1 for both fingers in the active condition and for the index finger in the passive condition. For the ring finger, the highest value in the passive condition was observed in BA 1.

**Figure 4.**
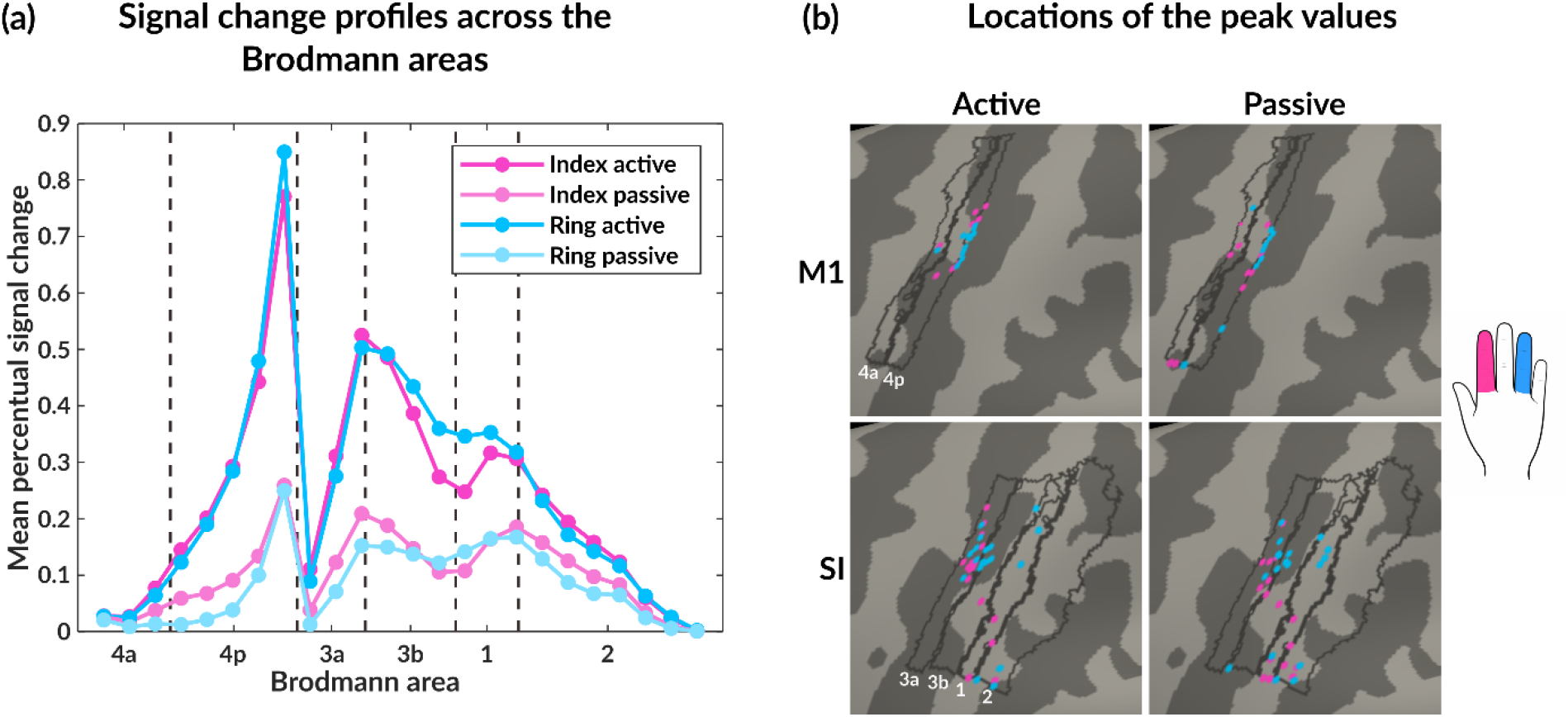
Signal change profiles and locations of the peak values. (a) Group-average percent signal change profiles across the Brodmann areas of the M1 and SI cortices for each finger and condition. Dashed vertical lines indicate the approximate boundaries between the Brodmann areas. (b) Vertices corresponding to the largest percent signal change for each finger and condition in the M1 and SI cortices. Index finger vertices are marked in pink and ring finger vertices are marked in blue. Boundaries of FreeSurfer’s probabilistic Brodmann areas are marked with gray lines.

### 3.2 Overlap and distance between fingers

Figures 3c and 3d show the comparison of the overlap of finger representations and the geodesic distance between fingers in the active and passive conditions in the M1 and SI cortices. To assess overlap between finger activations, a modified Dice coefficient between fingers was calculated for the passive and active conditions across both ROIs. Finger representations were more overlapping in the active than in the passive condition in the SI cortex (Supplementary table 1). In the M1 cortex, no statistically significant difference was observed. Finger separation was measured as the geodesic distance between the vertices corresponding to the largest percent signal change for each finger (Fig. 4b). No statistically significant difference in finger separation between the active and passive conditions was found in either ROI. However, there was a trend toward larger geodesic distances in the passive condition.

### 3.3 Subject-level activation patterns

Figure 5 shows the subject-level activation maps for each finger in the M1 and SI cortices for five representative participants (see Supplementary figures 1 and 2 for all of the maps). In the M1 cortex, passive movements elicited significant activation in 14 of 18 subjects for the index finger and in 13 of 18 for the ring finger, whereas active movements elicited activation in 17 of 18 subjects for both fingers. In the SI cortex, passive movements resulted in significant activation in 17 of 18 subjects, and active movements in all 18 subjects, for both fingers.

**Figure 5.**
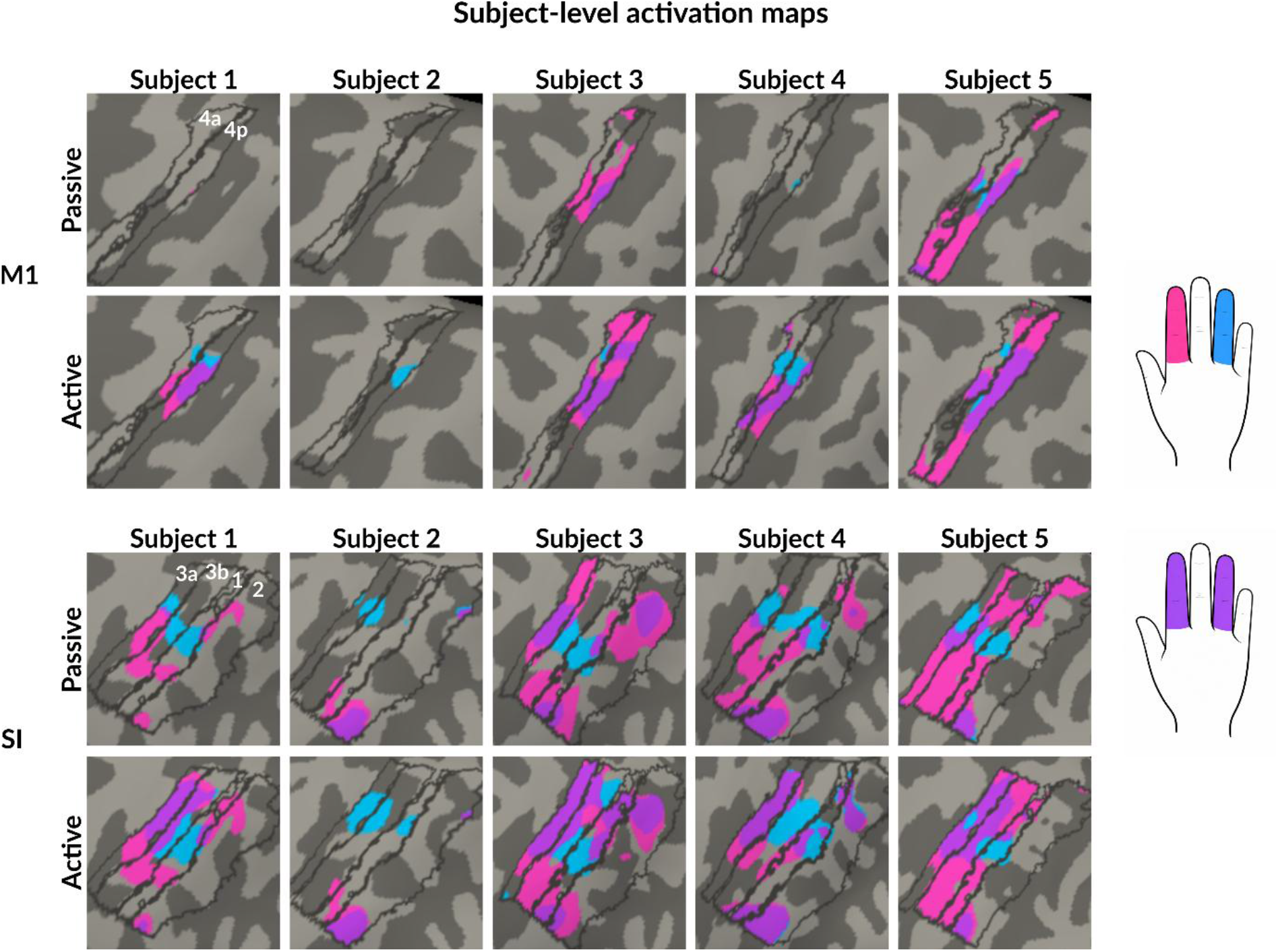
Subject-level activation maps. Activation maps for the index (pink) and ring (blue) fingers in the M1 and SI cortices for five representative subjects. Purple areas represent overlapping regions. Borders of FreeSurfer’s probabilistic Brodmann areas are marked with gray lines.

We also examined whether the finger representations followed somatotopy by calculating the average MNI x-coordinate from the finger-versus-rest maps. Table 1 shows the average MNI x-coordinate of the activations for each subject. If the representations followed a somatotopic organization, the index finger would be located more laterally, corresponding to a more negative MNI x-coordinate. In the active condition, somatotopic organization was observed in the M1 and SI cortices for 11 out of 16 and 11 out of 18 participants, respectively. In the passive condition, somatotopy was observed in 7 out of 13 participants in the M1 cortex and 7 out of 16 participants in the SI cortex.

**Table 1.** Somatotopic organization of the fingers. Average MNI x-coordinate of the activation for the index and ring fingers in the M1 and SI cortices for each subject and the average across all subjects. Coordinates that follow the expected somatotopic organization are shown in bold.

| Subject | M1 |  |  |  | SI |  |  |  |
| --- | --- | --- | --- | --- | --- | --- | --- | --- |
|  | Index passive | Ring passive | Index active | Ring active | Index passive | Ring passive | Index active | Ring active |
| 1 | **-34.96** | **-33.38** | **-35.31** | **-34.95** | **-42.22** | **-41.24** | -38.88 | -39.19 |
| 2 |  |  |  | -35.30 | **-49.98** | **-42.16** | **-50.43** | **-41.15** |
| 3 | -32.78 | -35.48 | -30.96 | -33.97 | -39.80 | -42.01 | **-39.28** | **-38.44** |
| 4 | **-45.51** | **-34.95** | -35.44 | -35.64 | -42.37 | -43.12 | -40.05 | -40.39 |
| 5 | -35.40 | -37.93 | -32.50 | -34.47 | -40.09 | -42.16 | **-39.58** | **-39.31** |
| 6 | -30.32 | -32.60 | -30.82 | -32.51 | -36.80 | -37.67 | -37.66 | -38.41 |
| 7 |  |  | -34.27 | -35.01 | -39.26 | -41.81 | -38.48 | -38.56 |
| 8 | **-35.05** | **-33.04** | **-34.81** | **-32.28** | **-40.78** | **-40.52** | **-39.61** | **-38.92** |
| 9 | **-35.44** | **-34.47** | **-33.06** | **-30.61** | -41.32 | -41.67 | **-40.01** | **-39.57** |
| 10 | **-33.78** | **-32.17** | **-33.10** | **-32.88** | **-38.12** | **-37.71** | -38.68 | -39.09 |
| 11 | **-31.86** | **-27.32** | **-31.58** | **-31.54** | **-39.70** | **-37.19** | -38.63 | -38.87 |
| 12 | -34.78 | -35.48 | **-34.53** | **-34.43** | -42.13 | -42.16 | **-41.65** | **-40.07** |
| 13 | **-34.27** | **-33.47** | **-32.51** | **-31.02** | -41.38 | -42.89 | **-40.46** | **-39.26** |
| 14 | -34.96 | -35.04 | **-34.79** | **-34.05** | **-43.18** | **-41.82** | **-41.05** | **-40.42** |
| 15 |  |  | -35.44 |  |  |  | **-39.58** | **-37.35** |
| 16 |  | -33.63 | **-31.99** | **-31.30** | -38.34 | -38.62 | **-36.48** | **-35.83** |
| 17 |  |  | **-36.22** | **-35.83** |  | -42.73 | -39.62 | -41.44 |
| 18 | -33.84 | -33.91 | **-34.80** | **-33.65** | **-40.13** | **-39.44** | **-40.61** | **-39.73** |
| Average | **-34.84** | **-33.79** | **-33.54** | **-33.38** | **-40.97** | **-40.76** | **-40.04** | **-39.22** |

### 3.4 Group-level activation maps

Figure 6a presents the group-level activation maps contrasting each condition against rest in the M1 and SI cortices. In the SI cortex, both the active and passive conditions produced significant activation for each finger. In contrast, the M1 cortex showed no significant activation during passive stimulation, while active movements elicited significant activation for both fingers. In the SI cortex, the activations were primarily located in BAs 3a, 3b, and 1. Overlapping regions were primarily located in BA 3a and BA 1 in the active condition, and in 3a in the passive condition.

**Figure 6.**
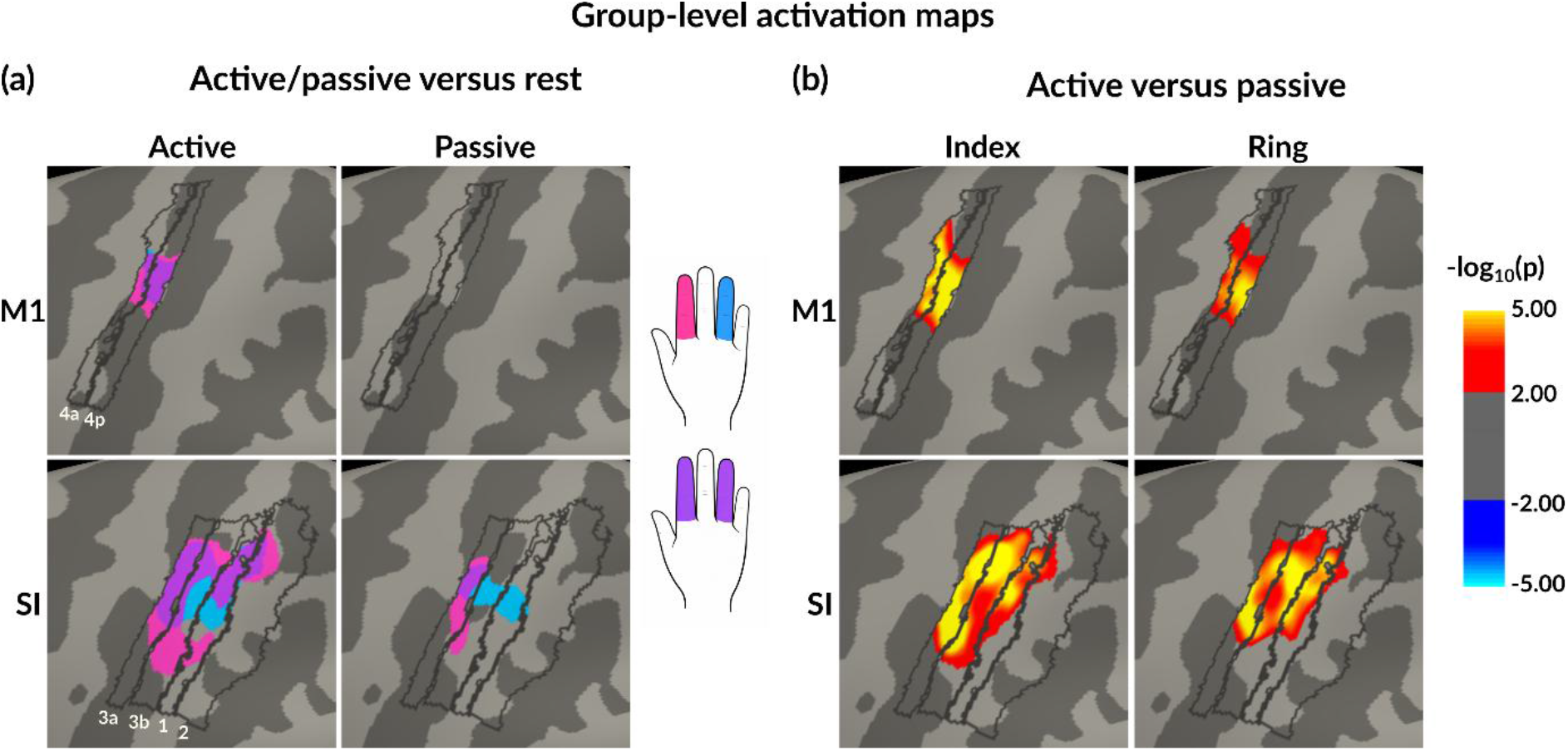
Group-level activation maps. (a) Maps contrasting the finger conditions (active/passive) against rest in the M1 and SI cortices. Significant activation for the index finger is marked in pink, for the ring finger in blue and the overlapping areas in purple. (b) Maps contrasting the active condition against the passive condition for both fingers in the M1 and SI cortices. The boundaries of the Brodmann areas are marked with gray lines.

Figure 6b presents the group-level activation maps contrasting active condition against passive condition for both fingers in the M1 and SI cortices. Activation in the active condition was stronger than in the passive condition in the same regions identified in the finger versus rest contrast. The passive condition did not show significantly stronger activation than the active condition in any ROI or for either finger.

## 4 Discussion

We investigated cortical representations of the index and ring fingers in the SM1 cortex during passive and active proprioceptive stimulation. The main finding was that the active condition elicited stronger and spatially more extensive BOLD responses than passive stimulation in both the SI and M1 cortices. The index and ring finger representations were more overlapping in the active than in the passive condition, but only in the SI cortex. However, finger representations were not significantly separate between the active and the passive conditions. At the individual level, activation patterns were more consistent across subjects in the active than in the passive condition, particularly in the M1 cortex. Together, these results suggest that volitional motor activation intensifies cortical processing of proprioceptive afference and modulates the organization of finger representations in the SM1 cortex, with a somatotopic organization present at the group level but variable across individuals.

### 4.1 Volitional finger activation intensifies cortical proprioceptive processing

The results indicated that SI and M1 cortical activations to proprioceptive stimulation are stronger and spatially more extensive in the active than in the passive condition for both fingers. The primary difference between the active and passive conditions is the active state of the SM1 cortex and the entire motor system involved required to maintain finger flexion, i.e. isometric type resistance, during the active condition, which may explain the stronger cortical activation through (1) facilitation of the SM1 cortical neurons, (2) intensified sensorimotor integration processing and (3) activation of neurons involved in the active motor control of the finger.

The stronger activation during the active than the passive condition was especially well demonstrated in the M1 cortex. Voluntary movement has been shown to increase the excitability of the M1 cortex, as demonstrated by previous transcranial magnetic stimulation (TMS) studies (Mazzocchio et al., 1994; Ugawa et al., 1995). In addition, although the finger movement was identical in both conditions, the active condition required cortical motor planning and motor output to “resist” the movement. In contrast, proprioceptive afference to the M1 cortex during the passive condition is likely suppressed or inhibited, potentially to prevent the execution of the sensed movement, analogous to previous magnetoencephalography (MEG) findings showing that inhibition suppresses motor output during viewing another person’s actions (Hari et al., 2014). In addition to the M1 cortex, stronger activation during the active than the passive condition was also observed in the SI cortex, which suggests that the processing of the proprioceptive afference may also be intensified during active tasks compared with passive ones. This may partly reflect the increased neuronal activity related to the intensified sensorimotor integration during the active condition (Thickbroom et al., 2003). A previous monkey study showed that during voluntary movements presynaptic inhibition of proprioceptive afferents at the level of the spinal cord is reduced by descending motor commands, thereby facilitating task-relevant proprioceptive input to spinal sensorimotor circuits (Tomatsu et al., 2023). However, it remains unclear to what extent such modulation at the spinal level directly influences the proprioceptive signals ascending to the cortex.

Proprioceptive signals originate from peripheral proprioceptors within the locomotor system that sense, e.g., the change in muscle length (muscle spindles) and muscle tension (Golgi tendon organs) (Proske & Gandevia, 2012). The muscle spindle is the only active proprioceptor receiving innervation to its intrafusal fibers through gamma-motoneurons during muscle activation, which allows the muscle spindle to be sensitive even when the muscle is contracting (i.e. shortening). Indeed, gamma-motoneuron input has shown to enhance proprioceptive afference even when the joint kinematics remain identical between conditions (Edin & Vallbo, 1988; Ribot-Ciscar et al., 1991). Therefore, it is possible that our observation of the stronger SM1 cortex response is, at least partly, due to increased peripheral stretch sensitivity of the muscle spindles during the active condition due to the gamma-motoneuron activation.

Finally, the stronger activation observed in the active condition may reflect not only enhanced proprioceptive processing, but also additional motor-related activity associated with voluntary muscle contraction. Because the experimental design did not include a condition involving resistance without proprioceptive stimulation, these contributions cannot be separated. The way activation strength was calculated may also have amplified the difference: average percent signal change was computed within a mask derived from all finger movement conditions versus rest, meaning that when the extent of activation was larger in the active condition, the average strength was likely higher as well.

Our findings align with previous studies demonstrating that proprioceptive processing is enhanced during an active motor task compared to passive movement. In nonhuman primates, proprioceptive signals have been shown to be amplified during active movements relative to kinematically matched passive movements already at the subcortical level in the cuneate nucleus (Versteeg et al., 2021). Similarly, in humans, an electroencephalography (EEG) study using corticokinematic coherence demonstrated that volitional muscle activation intensifies cortical proprioceptive processing in the SM1 cortex related to ankle-joint-movement stimulation (Giangrande et al., 2024). Although Giangrande et al. examined the foot and not the hand, their active condition closely resembled that of the present study: the active task required maintaining a static contraction rather than performing an active movement. In particular, the findings are in line with previous fMRI studies showing that active movements of the arm or fingers elicit stronger and more extensive BOLD responses than passive proprioceptive stimulation in the SM1 cortex (Lolli et al., 2019; Reddy et al., 2001; Thickbroom et al., 2003; Yu et al., 2011).

We also examined whether the group-average BOLD signal change differed across the Brodmann areas (Fig. 4a). In the M1 cortex, a peak in the percent signal change profile was observed in BA 4p in the active and passive conditions. BA 4p has been shown to be more strongly involved in the integration of sensory feedback during motor control than BA 4a, which is more closely associated with motor execution (Binkofski et al., 2002; Geyer et al., 1996). The predominant activation of BA 4p is therefore likely due to the proprioceptive input of the task, consistently observed in both passive and active conditions. In the active condition, however, this input is further integrated with active motor command, leading to increased sensorimotor integration. It is noteworthy to mention that in the active condition, peak activation in BA 4p was clearly stronger than the activation in the SI cortex. In the passive condition, the difference between the M1 and SI cortex activations was clearly smaller, which may be due to the absence of volitional motor output demand.

In the SI cortex, the strongest signal change peak was observed in BA 3a, close to the boundary between BA 3a and 3b for both fingers in the active condition and for the index finger in the passive condition (Fig. 4a). The corresponding peak for the ring finger in the passive condition was observed in BA 1. The peak in BA 3a was expected, as BA 3a is suggested to be the main cortical area for proprioceptive input (Iwamura et al., 1983; Krubitzer et al., 2004). The peak observed in BA 1 might be caused by some residual tactile stimulation, as BA 1 is typically associated with cutaneous processing (Purves, 2018). Nevertheless, it is important to note that the Brodmann area boundaries are only approximations both at the group and individual levels, and some degree of spatial variability or partial overlap between functional domains is likely also due to the vast reciprocal cortico-cortical connections between these areas.

### 4.2 Active condition produces more overlapping finger representations in the SI cortex

Finger representations overlapped more in the active than in the passive condition in the SI cortex. This observation may be because of overlaps in the functional anatomy. Individual fingers of the hand are used together, and they are anatomically linked by partially sharing the same parent muscles (Lang & Schieber, 2004). Thus, it seems that proprioceptive afference also activates overlapping neuronal networks in the SI cortex, and even more effectively during the active than the passive condition. However, the extent of the cortical activation to proprioceptive stimulus could also contribute to this effect. The more extensive activation in the active condition might result in greater overlap between fingers within the hand area. In the M1 cortex, a similar difference was not observed, which may be because the activation of the M1 cortex was weak in the passive condition, even below our detection threshold as only 12 subjects showed significant activation for both fingers in the passive condition.

The group-level activation maps indicated that the overlapping areas in the SI cortex were mainly located in BAs 3a and 1 in the active condition, whereas in the passive condition they were primarily located in BA 3a that is suggested to be the main area for proprioceptive processing (Iwamura et al., 1983; Krubitzer et al., 2004). This pattern aligns with the findings from nonhuman primates showing that, compared with the cutaneous area 3b, the topographic organization of area 3a is coarser, and the digit representations are less distinct and exclusive (Krubitzer et al., 2004).

The geodesic distance, reflecting the shortest path between the peak cortical signal change locations of the index and ring fingers, did not differ significantly when comparing the active and passive conditions. Nevertheless, it appeared that the fingers were more separated in the passive than in the active condition, which would have been in line with the more overlapping finger representations in the active than in the passive condition in the SI cortex.

### 4.3 Activation patterns vary across subjects and conditions

At the individual level, passive movements elicited significant activation for most of the participants in both the SI and M1 cortices, although it was more consistently observed across participants in the SI than in the M1 cortex. In the SI cortex, nearly all participants showed activation for both fingers (17 of 18), whereas in the M1 cortex the responses were more variable across individuals (14 of 18 for the index finger and 13 of 18 for the ring finger). At the group level, passive stimulation did not produce significant activation in the M1 cortex, suggesting some degree of inter-individual spatial variation in the M1 proprioceptive activation. Due to the absence of volitional motor output and lesser direct proprioceptive input to the M1 than to the SI cortex these observations were somewhat expected for the passive condition in the M1 cortex. In contrast, active movements elicited robust and consistent activation in both cortices, both at the individual (17 of 18 for the M1 cortex and 18 of 18 for the SI cortex) and group levels. This may reflect stronger cortico-cortical interaction between the SI and M1 cortices during the active condition.

Somatotopic organization was not observed in all participants. In the passive condition, it was observed in 7 of 13 participants for the M1 cortex and 7 of 16 for the SI cortex. In the active condition, it was observed in 11 of 16 participants for the M1 cortex and 11 of 18 for the SI cortex. This could be because the average MNI x-coordinate was calculated across the entire M1 and SI ROIs, which include several Brodmann areas that can show slightly different somatotopic layouts. However, at the group level, the average MNI x-coordinates followed the expected somatotopic organization, in line with previous group-level findings for tactile stimulation and active movements in the SI and M1 cortices (Dechent & Frahm, 2003; Martuzzi et al., 2012).

## 5 Conclusions

In conclusion, the results indicated that an active motor state of the SM1 cortex intensifies its cortical processing of proprioceptive afference from the hand and modulates the organization of the cortical finger proprioceptive representations when compared to identical proprioceptive stimulation in passive motor state. Proprioceptive finger representations followed the classical somatotopic organization at the group level, although this organization was not consistently observed across all subjects. Future research should extend these findings by examining all five fingers and by characterizing proprioceptive representations in greater detail across different Brodmann areas, and even beyond the SM1 cortex. In addition, further studies are needed to confirm cortical layer-specific patterns of the processing of proprioceptive afference during passive and active conditions.

## Acknowledgments

We would like to thank Jaana Hyyppö, Riikka Martin, and Satu-Maria Virtanen from Hospital Nova for their assistance with the measurements. We would also like to thank Markus Kerminen for building the proprioceptive stimulator.

