## Supplementary material for "Volitional hand activation intensifies cortical proprioceptive processing in the primary sensorimotor cortex"

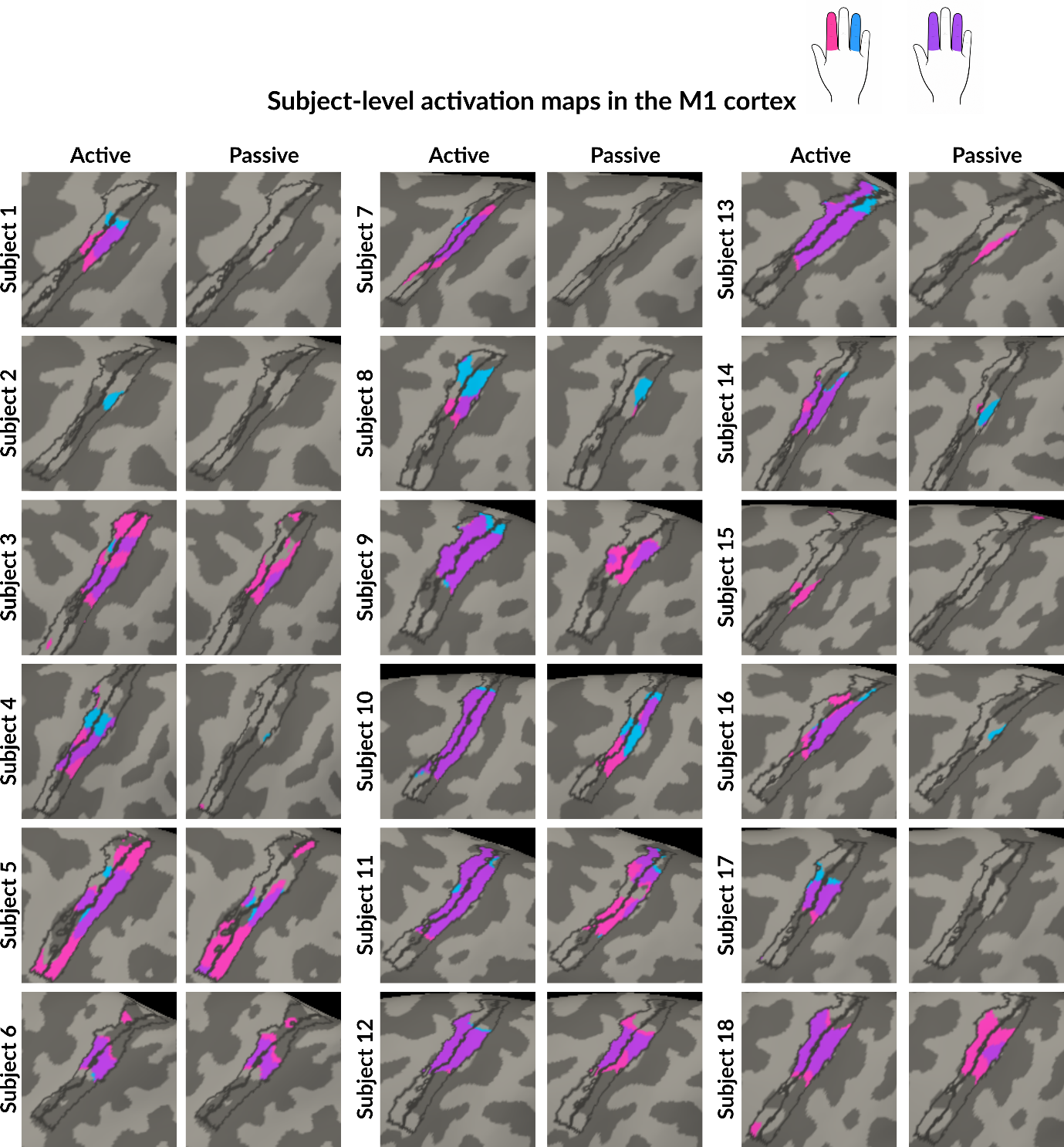
**Supplementary material**

**Supplemetary figure 1** Subject-level activation maps in the M1 cortex. Activation maps for the index (pink) and ring (blue) fingers are shown, each contrasted against rest. The maps are corrected for multiple comparisons using FDR at p < 0.05. Purple areas represent the overlapping areas. The boundaries of FreeSurfer’s probabilistic Brodmann areas are marked with gray lines.

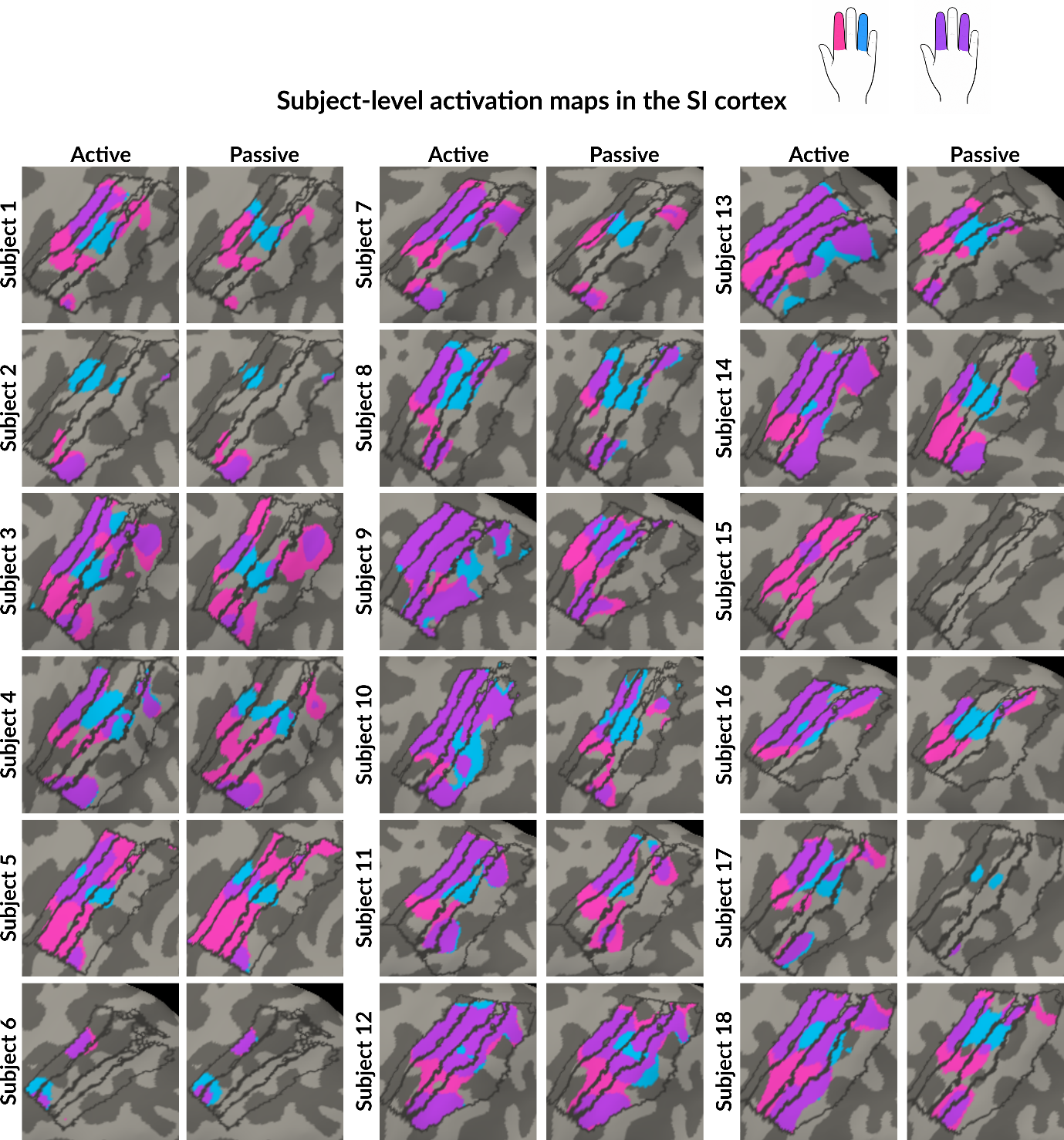
**Supplementary figure 2** Subject-level activation maps in the SI cortex. Activation maps for the index (pink) and ring (blue) fingers are shown, each contrasted against rest. The maps are corrected for multiple comparisons using FDR at p < 0.05. Purple areas represent the overlapping areas. The boundaries of FreeSurfer’s probabilistic Brodmann areas are marked with gray lines.

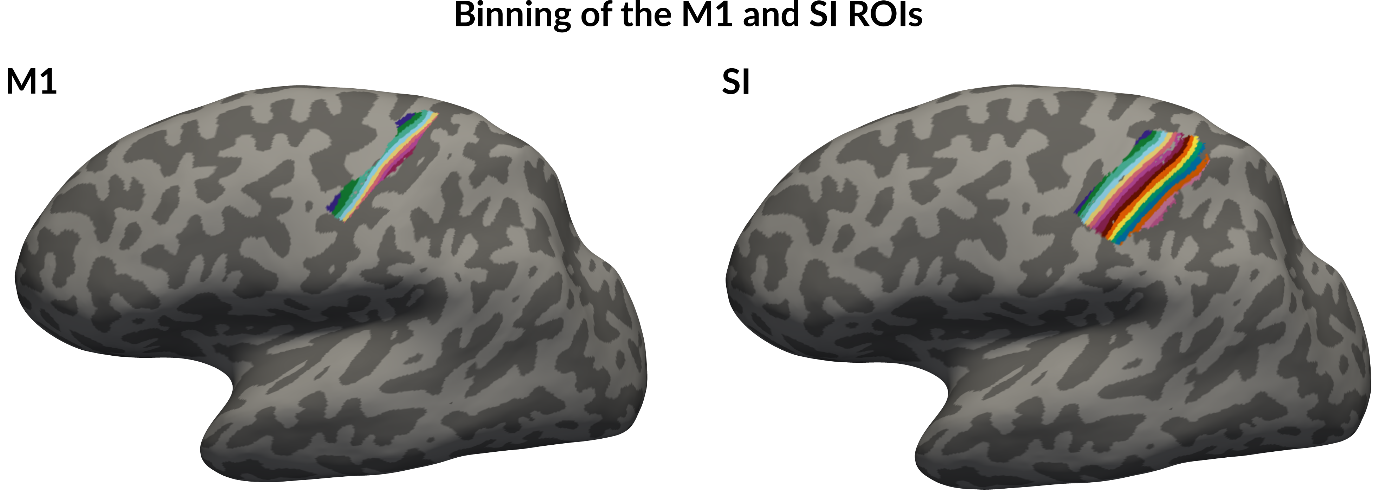
**Supplementary figure 3** Binning of the M1 and SI ROIs for signal profiles. The hand-area ROIs in the M1 and SI cortices were divided into 8 and 16 bins, respectively, to calculate the group-average percent signal change profiles across the Brodmann areas. To create the bins, a reference line was defined between two anatomically selected points within each ROI. For every vertex, the minimum Euclidean distance to this line was calculated. Vertices were then grouped according to these distances, resulting in a set of equally spaced bins across the ROI.

**Supplementry table 1** Comparison of metrics between the active and passive conditions (p-values and effect sizes). Statistically significant p-values are shown in bold.

|  | p-value | Effect size (r) |
| --- | --- | --- |
| Extent of activation compared between active and passive conditions |  |  |
| Index finger, M1 | **< 0.001** | 0.8808 |
| Ring finger, M1 | **< 0.001** | 0.8819 |
| Index finger, SI | **< 0.01** | 0.8209 |
| Ring finger, SI | **< 0.001** | 0.8783 |
| Strength of activation compared between active and passive conditions |  |  |
| Index finger, M1 | **< 0.001** | 0.8790 |
| Ring finger, M1 | **< 0.001** | 0.8790 |
| Index finger, SI | **< 0.001** | 0.8777 |
| Ring finger, SI | **< 0.001** | 0.8777 |
| Overlap of activations compared between active and passive conditions |  |  |
| M1 | 0.2661 | 0.3397 |
| SI | **< 0.05** | 0.6142 |
| Geodesic distance between fingers compared between active and passive conditions |  |  |
| M1 | 0.2631 | -0.3272 |
| SI | 0.0531 | -0.4880 |
